# Immortalised murine R349P desmin knock-in myotubes exhibit a reduced proton leak and decreased ADP/ATP translocase levels in purified mitochondria

**DOI:** 10.1101/2023.11.10.566522

**Authors:** Carolin Berwanger, Dominic Terres, Dominik Pesta, Britta Eggers, Katrin Marcus, Ilka Wittig, Rudolf J. Wiesner, Rolf Schröder, Christoph S. Clemen

**Author notes:** Both first authors contributed equally to this work (Carolin Berwanger, Dominic Terres). Author for correspondence: Christoph S. Clemen, Institute of Aerospace Medicine, German Aerospace Center (DLR), Linder Höhe, 51147 Cologne, Germany.

## Abstract

Desmin gene mutations cause myopathies and cardiomyopathies. Our previously characterised R349P desminopathy mice, which carry the ortholog of the common human desmin mutation R350P, showed marked alterations in mitochondrial morphology and function in muscle tissue. By isolating skeletal muscle myoblasts from offspring of R349P desminopathy and p53 knock-out mice, we established an immortalised cellular disease model. Heterozygous and homozygous R349P desmin knock-in and wild-type myoblasts could be well differentiated into multinucleated spontaneously contracting myotubes. The desminopathy myoblasts showed the characteristic disruption of the desmin cytoskeleton and desmin protein aggregation, and the desminopathy myotubes showed the characteristic myofibrillar irregularities. Long-term electrical pulse stimulation promoted myotube differentiation and markedly increased their spontaneous contraction rate. In both heterozygous and homozygous R349P desminopathy myotubes, this treatment restored a regular myofibrillar cross-striation pattern as seen in wild-type myotubes. High-resolution respirometry of mitochondria purified from myotubes by density gradient ultracentrifugation revealed normal oxidative phosphorylation capacity, but a significantly reduced proton leak in mitochondria from the homozygous R349P desmin knock-in cells. Consistent with a reduced proton flux across the inner mitochondrial membrane, our quantitative proteomic analysis of the purified mitochondria revealed significantly reduced levels of ADP/ATP translocases in the homozygous R349P desmin knock-in genotype. As this alteration was also detected in the soleus muscle of R349P desminopathy mice, which, in contrast to the mitochondria purified from cultured cells, showed a variety of other dysregulated mitochondrial proteins, we consider this finding to be an early step in the pathogenesis of secondary mitochondriopathy in desminopathy.

**Highlights:**

- R349P desminopathy immortalised murine myoblasts as a cellular disease model
- Electrical stimulation improves myofibrillar maturation in desminopathy myotubes
- Reduced proton leak in mitochondria of homozygous R349P desmin knock-in myotubes
- Reduced ADP/ATP translocase levels in mitochondria of desminopathy myotubes
- Early signs of secondary mitochondriopathy in desminopathy in cultured myotubes

## Introduction

Mutations in the human desmin gene (*DES*) cause familial and sporadic forms of desminopathies affecting skeletal and cardiac muscle tissue (OMIM #125660; (Brodehl et al., 2018; Clemen et al., 2013; van Spaendonck-Zwarts et al., 2010)). The human desmin mutation R350P (corresponding to R349P in mice) is the most common pathogenic desmin missense mutation in Germany (Walter et al., 2007). As muscle biopsy samples from desminopathy patients are scarce, patient-mimicking disease models are needed to study the sequential steps of the molecular pathogenesis. We had generated and characterised an R349P desmin knock-in mouse line that showed age-dependent skeletal muscle weakness, dilated cardiomyopathy and cardiac arrhythmias and conduction defects. In muscle cells, the intermediate filament protein desmin forms an extrasarcomeric cytoskeleton which connects myofibrillar Z-discs, intercalated discs, dense bodies, myonuclei, mitochondria, and the cell membrane (Clemen et al., 2013; Li et al., 1997). The R349P mutant desmin caused aberrant subcellular localisation and increased turnover of desmin and its binding partners, ultimately leading to disruption of the extrasarcomeric intermediate filament network (Clemen et al., 2015). In addition, we have previously described a marked secondary mitochondriopathy in human and murine desminopathy skeletal muscle, with focal depletion and accumulation of mitochondria and abnormally shaped mitochondria. At the molecular level, R349P mutant desmin led to a reduction in the amount and activity of complex I, a decrease in mtDNA-and ncDNA-encoded respiratory chain proteins, and prominent large-scale deletions in mtDNA (Winter et al., 2016). Furthermore, mitochondrial abnormalities were described in transgenic mice expressing L345P mutated desmin (Kostareva et al., 2008) as well as in desmin knock-out mice (Clemen et al., 2013; Elsnicova et al., 2022; Li et al., 1996; Li et al., 1997; Lindén et al., 2001; Milner et al., 2000; Milner et al., 1996; Thornell et al., 1997). Several previous studies of muscle biopsy specimens from human desminopathies and other forms of myofibrillar myopathies have also reported mitochondrial pathology (Clemen et al., 2013; Fidzianska et al., 2005; Jackson et al., 2015; Joshi et al., 2014; McCormick et al., 2015; Schröder et al., 2003, 2007; Vincent et al., 2016). To overcome the limitations of diagnostic muscle biopsies and to provide a tool to study the pathophysiology of desminopathies at the cellular level, we here generated improved cultures of immortalised myoblasts from soleus and gastrocnemius muscles of R349P desminopathy mice. In addition to a basic characterisation of this cellular disease model, we studied the mitochondrial pathology associated with the homozygous expression of R349P mutant desmin in more detail. This led to the identification of a reduced proton leak and decreased levels of ADP/ATP translocases in mitochondria purified from R349P desmin knock-in myotubes.

## Materials and methods

### Animals

Mice were housed in isolated ventilated cages (IVC) under specific and opportunistic pathogen-free (SOPF) conditions in a standard environment with free access to water and food. Health monitoring was done as recommended by the Federation of European Laboratory Animal Science Associations (FELASA). Mice were handled in accordance with the German Animal Welfare Act (Tierschutzgesetz) as well as the German Regulation for the protection of animals used for experimental or other scientific purposes (Tierschutz-Versuchstierverordnung). For muscle tissue dissection, mice were euthanized by cervical dislocation. All investigations were approved by the governmental office for animal care (Landesamt für Natur, Umwelt und Verbraucherschutz North Rhine-Westphalia (LANUV NRW), Recklinghausen, Germany (reference numbers 84-02.04.2014.A262 and addenda, 84-02.05.40.14.057, and 84-02.05.40.16.040).

### Generation of immortalised R349P desmin knock-in myoblasts from murine muscle tissue

Homozygous female R349P desmin knock-in mice (B6J.129Sv-*Des*^tm1.1Ccrs^/Cscl, MGI:5708562, (Clemen et al., 2015)) were crossbred with homozygous male p53 knock-out mice (B6J.129S2-*Trp53*^tm1Tyj^/J, MGI:1857263, (Jacks et al., 1994)), and double-heterozygous offspring was further mated to obtain mice with the following allele combinations: i) homozygous R349P desmin knock-in and homozygous p53 knock-out, ii) heterozygous R349P desmin knock-in and homozygous p53 knock-out, and iii) desmin wild-type and homozygous p53 knock-out. These mice were used for the isolation of myoblasts separately derived from the soleus and gastrocnemius muscles. The obtained myoblasts are immortalised due to the lack of p53, as described earlier (Jacks et al., 1994; Metz et al., 1995; Winter et al., 2019). Though crossing-in the p53 allele (Jacks et al., 1994) is a time-consuming approach, it has the advantage of being a targeted immortalization technique.

The isolation of skeletal myoblasts was performed following published methods (Rando and Blau, 1994; Rosenblatt et al., 1995; Winter et al., 2019) with modifications. Mice at 2 to 3 weeks of age were euthanized and soleus and gastrocnemius muscles were dissected separately and enzymatically digested (per mouse both soleus muscles in one tube, and both gastrocnemius muscles in another tube) in 4 mL preparation buffer (0.2% collagenase I in DMEM with 1 mM pyruvate, 1x non-essential ammino acids, 2 mM L-glutamate, 100 U/mL penicillin and 0.1 mg/mL streptomycin) by shaking at 200 rpm for 2h at 37°C. Digested tissues were centrifuged at 200x g for 5 min, resuspended in plating buffer (DMEM, 20% fetal calf serum, 10% horse serum, 1% chicken embryo extract, 1 mM pyruvate, 1x non-essential ammino acids, 2 mM L-glutamate, 100 U/mL penicillin and 0.1 mg/mL streptomycin) and transferred into a 100 mm diameter cell culture dish (# 83.3902, Sarstedt, Nümbrecht, Germany). The partially dissociated, digested muscles were further separated into single muscle fibers by carefully pipetting (10 mL plastic cell culture pipette) the suspension, plated on a collagen-coated (0.2% collagen for 1h at 37°C) 100 mm diameter cell culture dish, and incubated at 37°C and 5% CO_2_ for 6 to 7 days until satellite cells had emerged from the muscle fibers on the coated cell culture dish. After two to three days, fresh plating medium was added to the cells.

For quality control, the desmin and p53 genotypes of each dissected mouse were verified by PCR genotyping. Primers for the R349P desmin knock-in allele were 5’-AAACCTGGAAGCAGTTTTACACAAGAGGC-3’ and 5’-GCTGTAGGTTTTTAATTCTAAAGGTGGATAAGGG-3’ (Clemen et al., 2015) and for the p53 knock-out allele 5’-TATACTGAGAGCCGGCCT-3’, 5’-ACAGCGTGGTGGTACCTTAT-3’, and 5’-CATTCAGGACATAGCGTTGG-3’ (Jacks et al., 1994). Furthermore, each established culture of myoblasts was verified by desmin and p53 PCRs and by desmin immunofluorescence analysis. The immortalised myoblast cultures used for this work had passage numbers of maximal 35, and from each culture used for experiments the desmin genotype was repeatedly verified.

Notes of caution: Every immortalised myoblast ‘line’ is derived from either the soleus muscles or the gastrocnemius muscles of one individual mouse so that the derived myoblast cultures are polyclonal. With increasing passage, the number of differentiated myotubes may decrease. Almost all myoblast cultures contained a fraction of unwanted fibroblasts which need to be controlled as described below. Such fibroblast contamination cannot be eliminated by addition of choleratoxin, as it was shown to induce myoblast fusion (Stygall and Mirsky, 1980). Importantly, the myoblasts do neither grow on glass surfaces nor on treated glass surfaces like on collagen or poly-L-lysine coated or on plasma-treated cover slips.

### Cultivation of myoblasts and reduction of ‘contaminating’ fibroblasts

For further cultivation myoblasts were split at a maximum of 70% confluence. Myoblasts were washed with 1x PBS and trypsinised using 0.05% trypsin/0.5 mM EDTA (Pan Biotech, Aidenbach, Germany). Trypsin was then stopped with Trypsin Neutralizing Solution (TNS, C-4111, Promocell, Heidelberg, Germany), and cells were centrifuged at 500x g for 8 min. Subsequently, pellets were resuspended in Skeletal Muscle Cell Growth Medium with supplement (SkMCG-Medium C-23060 (contains basal medium C-23260B and supplement mix C-39365), Promocell, Heidelberg, Germany) and with 100 U/mL penicillin and 0.1 mg/mL streptomycin included. The resuspended myoblasts were preplated on uncoated cell culture dishes for 1h to separate unwanted fibroblasts from the myoblasts.

After this time, most of the ‘contaminating’ fibroblasts, if present, were attached to the uncoated plastic dish, and the supernatant containing the myoblasts was diluted 1:1 to 1:10, depending on the amount, and transferred to 0.2% collagen-coated dishes for further cultivation.

In addition to preplating, we purified some of the myoblast cultures using the mouse Satellite Cell Isolation Kit (130-104-268, Miltenyi Biotec, Bergisch-Gladbach, Germany) when necessary due to higher fractions of fibroblasts (e.g., more than 20%). After trypsinisation and centrifugation, cells were resuspended in 100 µL SkMCG-Medium, and 20 µL magnetic beads were added and the suspension was incubated for 15 min at 4°C. Myoblasts (like satellite cells for which this kit was intended) did not bind to the magnetic beads, which were coated with multiple antibodies directed against non-muscle cell surface antigens, while other unwanted cells including the fibroblasts bound to the beads. MS columns (130-042-201, Miltenyi Biotec, Bergisch-Gladbach, Germany) were placed in the OctoMACS Separator (Miltenyi Biotec, Bergisch-Gladbach, Germany) and pre-washed with SkMCG-Medium. Subsequently, 0.2% collagen coated dishes already containing SkMCG-Medium were placed below the columns. 500 µL medium were pipetted to the cells-beads-mixture which was added to the MS column, and the myoblasts flowed through the column and were collected in the culture dish. Additional 500 µL medium were used to rinse the columns twice.

### Differentiation of myoblasts into mature myotubes

The immortalised myoblasts were differentiated into myotubes at approximately 90% confluence with a standard differentiation medium (DMEM, 5% horse serum, 1 mM pyruvate, 1x non-essential ammino acids, 2 mM L-glutamate, 100 U/mL penicillin and 0.1 mg/mL streptomycin) for 5 to 10 days. Every 2 to 3 days half of the differentiation medium was replaced with fresh one. This yielded a significant number of multinucleated myotubes.

To generate more mature myotubes, the myoblast cultures were differentiated by adding differentiation medium in combination with electrical pulse stimulation for 4 to 7 days. Electrical pulse stimulation was performed according to (Grande et al., 2023; Orfanos et al., 2016) with modifications; the settings for this long-term stimulation (‘promoting protocol’) were 0.05 Hz, 4 ms, 10 V (Reimann et al., 2020) using a C-Pace EM Tissue Culture Interface equipped with Stretch/High Voltage Driver Boards and 6-dish C-Dish Culture Dish Electrode Systems (TCI100, SHV100, CLD6W35CN; Ion Optix, Milton, MA, USA).

### Microscopic imaging

Phase contrast microscopic images of myoblasts and myotubes grown in 100 mm diameter culture dishes were recorded with a transmission light microscope (DMIL LED; Leica Microsystems, Wetzlar, Germany), objectives N PLAN 5x/0.12 PH0 and HI PLAN I 10x/0.22 PH1, equipped with a heated stage (Tempcontrol-37; Pecon, Erbach, Germany) and stand-alone FLEXACAM C1 colour camera (Leica Microsystems, Wetzlar, Germany).

For immunofluorescence analysis, myoblasts and myotubes grown in 35 mm plastic µ-dishes (#81156, Ibidi) were fixed in 4% paraformaldehyde in PBS at RT for 20 min, permeabilised with 0.5% Triton X- 100 in PBS for 10 min, incubated in 0.15% glycine in PBS for 10 min to block residual formaldehyde, and non-specific binding sites were blocked with 1% BSA at RT for 1h. Cells were incubated with primary antibodies diluted in 0.5% BSA in PBS at 4°C overnight in a humid chamber. Secondary antibodies and DAPI were diluted in 0.5% BSA in PBS and added for 1h at RT. Each step was followed by several washings in PBS and finally cells were rinsed once with ddH_2_O and embedded in Mowiol/DABCO. Confocal images were recorded using an Infinity Line system (Abberior Instruments GmbH, Göttingen, Germany), the UPLXAPO60XO NA 1.42 objective, and software Imspector version 16.3.16100 in LightBox mode. Images were deconvolved using Huygens Professional version 23.04.0- p2 (Scientific Volume Imaging B.V., Hilversum, The Netherlands). Primary antibodies used were directed against desmin (mouse monoclonal, M0760, Agilent/Dako, 1:100), α-actinin-2 (mouse monoclonal, A7811, Sigma, 1:200), and VDAC1 (rabbit polyclonal, ab15895, Abcam, 1:200). Secondary antibodies used were either goat anti-mouse IgG (H+L) cross-adsorbed Alexa Fluor 488 (A-11001) or goat anti-mouse IgG (H+L) cross-adsorbed Alexa Fluor 568 (A-11004) combined with goat anti-rabbit IgG (H+L) cross-adsorbed Alexa Fluor 647 (A-21244; all from Thermo Fisher, 1:400).

### Purification of mitochondria by density gradient ultracentrifugation

Per 13 mL ultracentrifugation tube, a total number of four 150 mm diameter cell culture dishes containing well-differentiated myotubes of either wild-type or homozygous R349P desmin knock-in genotype were harvested using a rubber policeman. Harvested cells were washed with PBS twice and centrifuged at 1,500x g for 15 min at RT. All subsequent steps were performed at 4°C without loss of time to ensure a high coupling state of the mitochondria for respirometry. The cell pellet was resuspended in 1.2 mL ice-cold hypo-osmotic cell homogenisation medium with protease inhibitors (CHMP: 150 mM MgCl_2_, 10 mM KCl, 10 mM Na-succinic acid, 1 mM ADP potassium salt, 0.25 mM dithiothreitol (DTT), 10 mM Tris/HCl, Roche Complete Mini w/o EDTA, 2 mg/mL Pepstatin A, pH 6.7). Mg^2+^ and K^+^ were added to protect nuclei and to prevent cytoplasmic protein gel formation. Succinic acid, ADP and DTT as well as sucrose (see below) were added to support functional and mechanical integrity of the mitochondria. Resuspended cells were homogenised in a 5 mL Dounce homogeniser (Wheaton Type B) by ten quick strokes of a tight-fitting puncher avoiding foam formation, and cell disruption was confirmed by phase contrast microscopy. The homogenate was mixed with 400 µL CHMPSa (CHMP containing 1 M sucrose) and centrifuged for 10 min at 1,500x g to remove nuclei and unbroken cells. The supernatant was collected, and the pellet was resuspended in 1 mL CHMPSb (CHMP containing 0.25 M sucrose). This suspension was again homogenised by ten strong strokes of a tight-fitting Dounce homogeniser and centrifuged for 10 min at 1,500x g. This step was repeated until three supernatants were collected; the final nuclear pellet was discarded. Subsequently, the supernatants were centrifuged for 15 min at 15,000x g to collect crude mitochondrial fractions. The pellets were collected and combined in 1.2 mL ice-cold iodixanol buffer A (IBA: 0.25 M sucrose, 1 mM Na_2_EDTA, 10 mM Na-succinic acid, 1 mM ATP potassium salt, 0.25 mM DTT, 10 mM HEPES, pH 7.4) and resuspended by five gentle strokes of a tight-fitting Dounce homogeniser. At last, 1.8 mL of a 50% iodixanol buffer solution prepared from Optiprep (D1556, Sigma-Aldrich) and IBA was added to give a final iodixanol concentration of 30%.

A 9-step discontinuous isosmotic iodixanol density gradient was prepared in a 13.2 mL open-top thinwall ultra-clear centrifugation tube (#344059, Beckman Coulter) using Optiprep and iodixanol buffer B (IBB: 0.25 M sucrose, 6 mM Na_2_EDTA, 60 mM HEPES, pH 7.4). The gradient contained the following density steps of iodixanol, 23%, 21%, 19%, 17%, 15%, 13%, 11%, 9% and 7%, spanning a density range of approximately 1.10-1.18 g/mL. Each density gradient tube was allowed to equilibrate for 4 h at 4°C. The gradient was underlaid with the above crude mitochondrial preparation in 30% iodixanol using a 5 mL syringe and an 80 mm long metal cannula (#155456G21, B. Braun, Melsungen, Germany) carefully avoiding movement of the gradient, e.g., by air bubbles. Finally, the gradient was overlaid up to maximum filling height with IBA (Fig. 4A), tared and ultracentrifuged for 1 h at 110,000x g using a Beckman L7-65 ultracentrifuge with a SW41Ti rotor (#331362, Beckman Coulter). After ultracentrifugation, the gradient tube showed several distinct white-turbid regions enriched in mitochondria (Fig. 4B). The central region comprising three white-turbid bands was collected and centrifuged for 20 min at 20,000x g. The pellet containing the isolated mitochondria was immediately resuspended in 100 µL ice cold mitochondrial respiration medium (MiR05 according to https://mitoglobal.org/images/c/c8/MiPNet22.10_MiR05-Kit.pdf: 0.5 mM EGTA, 3 mM MgCl_2_, 60 mM lactobionic acid, 20 mM taurin, 10 mM KH_2_PO_4_, 20 mM HEPES, 110 mM sucrose, 1 g/l BSA fatty acid free, pH 7.1) for subsequent respirometry and immunoblotting. For the latter, an aliquot of 20 µL was mixed 1:2 to 1:5 (depending on the turbidity of the solution of resuspended mitochondria) with 5x SDS-PAGE sample buffer.

### High-resolution respirometry of purified mitochondria

Respirometry was performed using two two-chambered Oxygraph 2k devices equipped with DatLab software v.7.4.0.4 (Oroboros Instruments, Innsbruck, Austria). Both instruments were operated in parallel and were used to measure wild-type mitochondria in one and desminopathy mitochondria in the other chamber of the same device. Each respirometer chamber contained 2 mL MiR05 buffer at 37°C, air calibration was performed, and background oxygen flux was validated between 2 to 4 pmol s^−1^ mL^−1^ to rule out biological contamination (Pesta and Gnaiger, 2012). Volumes of 20 µL of either the above wild-type or desminopathy mitochondria suspensions were injected into the chambers. Two different substrate-uncoupler-inhibitor titration (SUIT) protocols were performed; in one device complex I-and in the other device complex II-dependent respiration of wild-type and desminopathy mitochondria was measured. Equilibrium was waited for between injections into the chambers. Oxygen levels in the chambers were maintained between 50 to 250 µM to prevent the limitation of oxidative phosphorylation by insufficient oxygen concentrations (cO_2_). At a cO_2_ below 50 µM the experiment was paused, the chamber was slightly opened, and the cO_2_ was allowed to rise above 200 µM again. The respiratory coupling control states were as follows: i) a non-phosphorylating resting state in the absence of adenylates, called LEAK state (L_N_) or state 4 in isolated mitochondria, ii) an ATP-generating oxidative phosphorylation with oxygen flux attributable to highly coupled active respiration, called OXPHOS state or state 3 in isolated mitochondria, iii) a maximally uncoupled respiration induced by protonophores such as FCCP, called ET-capacity or state 3u in isolated mitochondria, and iv) a residual oxygen consumption after depletion of substrates, oxygen, or total inhibition of electron transfer, called ROX state or state 2. All states were corrected for ROX.

In the chambers of one device, the substrate-uncoupler-inhibitor titration protocol SUIT 012 (https://wiki.oroboros.at/index.php/SUIT-012_O2_mt_D027) was performed. Only NADH pathway (N-pathway) stimulating substrates (final concentrations) were added, pyruvate (5 mM) and malate (2 mM) to achieve L_N_. At this point, respiration mostly compensated for proton leak and proton slip across the inner mitochondrial membrane. Active respiration was stimulated by addition of saturating levels of ADP (5 mM) resulting in OXPHOS state. In the absence of succinate, no significant stimulation by complex II could be expected. Cytochrome c (10 µM) was added to test the integrity of the outer mitochondrial membrane (_mt_OM) indicated by fractional O_2_ flux stimulation and compensated for potential cytochrome c loss that would otherwise limit OXPHOS state. Addition of yet another N-pathway substrate glutamate (10 mM) induced maximum complex I-linked OXPHOS capacity (C_I_P). By titration of the uncoupler carbonyl cyanide p-trifluoro-methoxyphenyl hydrazone (FCCP; 0.25 µM steps), the non-coupled electron transfer capacity of complex I was examined (C_I_E). An equilibrated maximum uncoupled ET-state was achieved at FCCP concentration of around 2-3 µM. Finally, complex III inhibitor antimycin A (2.5 µM) was injected to prevent any electron flow through Q-enzyme. Total inhibition of the electron transfer pathway allowed for the measurement of ROX. In the chambers of the other device, the substrate-uncoupler-inhibitor titration protocol SUIT 020 (https://wiki.oroboros.at/index.php/SUIT-020_O2_mt_D032) was performed to assess the Q-junction additivity of mostly complex I (N-pathway) and complex II (S-pathway). Pyruvate (5 mM) and malate (2 mM) were added to achieve L_N_. ADP, cytochrome C and glutamate were added to initiate the OXPHOS state and to check for _mt_OM integrity. Addition of succinate (20 mM) induced maximum OXPHOS capacity supported by joint activity of the NS-pathway (CI+II_P_). The convergent electron flow to the Q-junction from NADH dehydrogenase at complex I and succinate dehydrogenase at complex II reconstituted the functionality of the TCA cycle in intact mitochondria. Further, addition of rotenone (5 µM) inhibited complex I activity as well as residual electron flow from the fatty acid oxidation pathway. Remaining O_2_ flux was therefore attributable to complex II-activity (CII_P_). Oligomycin (1 µM) was added to induce a non-phosphorylating LEAK respiration, or state 4, by inhibiting ATP synthase. Titration of FCCP resulted in maximum uncoupled ET capacity. The total inhibition of electron transfer pathway was induced by inhibiting complex III via the addition of antimycin A (2.5 µM) resulting in ROX state.

### Immunoblotting and densitometric analysis

For Western blotting, aliquots of the isolated mitochondria were lysed in 5x SDS-PAGE sample buffer (125 mM Tris-HCl pH 6.8, 4% SDS, 20% glycerol, 10% 2-mercaptoethanol, 0.005% bromophenol blue; (Laemmli, 1970)), and proteins were separated by 15% SDS-PAGE and transferred to 0.2 µm Nitrocellulose Blotting Membrane (#10600001, GE Healthcare Life Sciences) by the semi-dry method (Towbin et al., 1979). The membranes were incubated with the fluorescent No-Stain Protein Labeling Reagent (A44449, ThermoFisher Scientific) according to the manufacturers protocol to detect total proteins using an ECL ChemoStar Imager HR 9.0 (Intas Science Imaging) in fluorescence mode (blue LED light source (470/30 nm), red emission filter (716/40 nm), exposure time 30 s). Subsequently, membranes were blocked with Tris-buffered saline-Tween-20 (TBS-T) buffer (10 mM Tris-HCl pH 8.0, 150 mM NaCl, 0.2% Tween-20) containing 5% milk powder. Primary antibodies were diluted in TBS-T buffer and incubated overnight; secondary antibodies coupled to horseradish peroxidase (POD) were diluted in TBS-T and incubated at RT for 1h. Primary antibodies used were directed against VDAC1 (rabbit polyclonal, ab15895, Abcam, 1:1,000), citrate synthase (CS) (mouse monoclonal, MA5-17264, Invitrogen, 1:3,000), and TOMM20 (mouse monoclonal, H00009804-M01, Abnova, 1:250). Secondary antibodies were anti-mouse or anti-rabbit-POD (A4416 and A6154, respectively, Sigma-Aldrich, 1:10,000). Visualisation was done by enhanced chemiluminescence using the SuperSignal West Pico PLUS chemiluminescent substrate (#34577, Thermo Scientific) and the ECL ChemoStar Imager HR 9.0 (Intas Science Imaging) in Chemiluminescence mode. After protein transfer, SDS-PA gels were stained with Coomassie Brilliant Blue, and remaining, non-transferred protein content was imaged i) using a white lightplate (Kaiser slimlite plano, #2453, 5,000 K) and the ECL ChemoStar Imager HR 9.0 (Intas Science Imaging) in transillumination mode as well as ii) the imager in ‘Coomassie’ fluorescence mode (red LED light source (628/35 nm), red emission (716/40 nm) filter, exposure time 60 s). The software LabImage 1D version 4.2.1 (Kapelan Bio-Imaging GmbH, Leipzig, Germany) was used for quantitation of the total protein per line signals obtained by No-Stain fluorescence and Coomassie transillumination.

### Analysis of high-resolution respirometry data

The respirometry results were visualised graphically online to provide a comprehensive view of oxygen consumption over time. Using the DatLab software version 7.4.0.4 (Oroboros Instruments GmbH, Innsbruck, Austria), curve proportions of the different measurement sections with a steady state condition between injections were chosen to compute the average oxygen fluxes and their derivatives. The data obtained were then related to the total protein content of the mitochondrial suspensions injected into the respirometer chambers. The factors used for these adjustments were calculated from the dilution factors of the SDS-PAGE sample preparations and loading volumes, and the normalised averages of the No-Stain fluorescence-and Coomassie Brilliant Blue absorbance-based line densitometry values. The data transformed in this way were respirometry values specific to the mitochondrial amounts measured. They were imported into GraphPad Prism version 9.4.1 (GraphPad Software, Boston, MA) in grouped tables and the values of the different measurement sections were statistically analysed using the non-parametric Mann-Whitney U (Wilcoxon rank-sum) test without multiple testing correction. The number of experiments and significance levels for each analysis are indicated in the Results section.

### Proteomic analysis of purified mitochondria

Aliquots of 20 µL of the samples of purified mitochondria lysed in 5x SDS-PAGE sample buffer for Western blotting were used for SDS-PAGE and tryptic in-gel digestion as described (Plum et al., 2020). Peptide concentration was determined via amino acid analysis (Eggers et al., 2021), and 200 ng of peptides were used for mass spectrometric measurements. Liquid-chromatography tandem mass spectrometry (LC-MS/MS) analysis, was carried out as described (Wulf et al., 2022) with modifications. In brief, LC-MS/MS measurements were performed on an Ultimate 3000 RSLC nano LC system (Dionex, Idstein, Germany) coupled to an Orbitrap Fusion Lumos Tribrid mass spectrometer (Thermo Fisher Scientific). The LC system was operated with a pre-column with a flow rate of 30 mL/min (Acclaim PepMap nanoViper, Thermo Fisher Scientific; 100 µm x 2 cm, 5 µm particle size) and an analytical C18 column (Acclaim PepMap nanoViper, Thermo Fisher Scientific; 75 µm x 50 cm, 2 µm particle size) with a flow rate of 400 nL/min. Separation of peptides was performed utilising a gradient starting with 95% solution A (0.1% FA) and 5% solution B (84% acetonitrile, 0.1% FA) with increasing solution B up to 30%. For DDA, a scan range from 350 to 1,400 m/z, a resolution of 60,000 and a maximum injection time of 50 ms were used. The Top15 selected precursors were fragmented with a fixed collision energy of 32% by higher-energy collisional dissociation (HCD) and a dynamic exclusion for 80 s. Fragment ion scans were performed at a resolution of 30,000 with a maximum injection time of 120 ms. Raw data of the purified mitochondria analysis was deposited in the PRIDE repository under the identifier PXD044624.

The MaxQuant software (Cox and Mann, 2008) version 2.0.3.0 was used for the analysis of raw files. The resulting peak lists were searched against the *mus musculus* UniProt FASTA reference proteome version (download 03/2023) and a common contaminants database provided in the Andromeda search engine (Cox et al., 2011; Tyanova et al., 2016a). Oxidation of methionine was set as variable modification and trypsin as digestion enzyme (maximum number of two missed cleavages). The false discovery rate (FDR) was set to 1% for peptides (minimum length of seven amino acids) and proteins. Label free quantification (LFQ) was performed using the MaxQuant LFQ normalisation whereby unique and razor peptides were assessed for protein quantification (Cox et al., 2014). Additionally, intensity based absolute quantification (iBAQ) value calculation was enabled. iBAQ values were normalised for each sample and adjusted to *per mille* values. Statistical analysis was carried out in the Perseus software (Tyanova et al., 2016b) version 2.0.3.0. Contaminants and decoys were filtered out and LFQ values were log_2_ transformed for further statistical analysis. A minimum of 70% valid values in at least one group was set as an additional filtering criterion prior to relative quantification. Remaining missing values were imputed from a normal distribution using a width of 0.3 and a downshift of 1.8. Paired Student’s t-test as well as permutation-based FDR correction were applied to the dataset (q-value) to determine statistically significant differential proteins.

### Proteomic analysis of soleus muscle tissue

Soleus muscles from five mice per genotype were pooled, and lysates prepared from this material of heterozygous and homozygous R349P desmin knock-in mice and wild-type littermates were subjected to label-free quantitative mass spectrometry analysis as described in detail in (Winter et al., 2016). The full data set of the soleus muscle proteomic analysis from this previous study has been re-assessed for the purposes of this work. In brief, MS data were analysed using MaxQuant version 2.0.1.0 (Cox and Mann, 2008). Enzyme specificity was set to trypsin, variable modifications were N-terminal acetylation, oxidation of methionine, and fixed modification was carbamidomethylation of cysteines. The option ‘matched between runs’ was enabled. The mouse reference proteome database (downloaded from Uniprot, August 26^th^ 2023, 63,361 entries, canonical and isoform entries) was used to identify peptides and proteins. The false discovery rate (FDR) for proteins and peptides was 1%. Identified and quantified proteins were further analysed using Perseus version 1.6.1.3 (Tyanova et al., 2016b). Label-free quantification values were normalised to the median of each sample. Raw data has now been deposited in the PRIDE repository under the identifier PXD046706.

### Miscellaneous methods

Respirometry graphs were generated using GraphPad Prism version 9.4.1 (GraphPad Software, Boston, MA), proteomics graphs using Excel 2016 (Microsoft) in conjunction with the add in “XY Chart Labeler” version 7.1 by Rob Bovey available at http://www.appspro.com/ and the add-in „Real Statistics Resource Pack” release 8.7 by Charles Zaiontz available at http://www.real-statistics.com. Graphs and images were further processed and figures assembled using CorelDraw Graphics Suite X7 (Corel Corporation, Ottawa, Canada).

## Results

### Long-term electrical pulse stimulation promotes myofibril organisation in immortalised myotubes expressing mutant desmin

Building on our previous work on the generation of immortalised skeletal muscle myoblasts carrying the R349P desmin mutation (Winter et al., 2019), we generated new cultures of immortalised myoblasts from either soleus or gastrocnemius muscles of heterozygous and homozygous R349P desmin knock-in mice and wild-type siblings using a more advanced experimental approach. Integrated modulation contrast (IMC) microscopy images showed myoblasts with round to spindle-shaped morphology, essentially free of contaminating fibroblasts, and no obvious differences between genotypes (Fig. 1, top row). Desmin immunofluorescence images showed a normal intermediate filament network in wild-type myoblasts, in contrast to homozygous cells, which displayed a severe disruption of the desmin intermediate filament network with desmin aggregates and few filament fragments. Heterozygous myoblasts exhibited a mixed pattern in the form of desmin filaments and aggregates that varied from cell to cell (Fig. 2, left column). VDAC1 labelling of mitochondria showed a distribution that was indistinguishable between wild-type and heterozygous cells, whereas mitochondria in homozygous myoblasts were present in granular structures with increased VDAC1 signal intensity (Fig. 2, middle column). Differentiation of myoblasts using standard differentiation medium resulted in the formation of myotubes (Fig. 1, 2^nd^ row) which occasionally showed spontaneous contractions in all three genotypes. While continuous electrical pulse stimulation for one week had no apparent effect on myotube morphology (Fig. 1, 3^rd^ row), this treatment markedly increased the number of spontaneously contracting myotubes. Visualisation of myofibrils using α- actinin-2 antibody staining showed a regular cross-striation pattern, indicating the presence of multiple, well-aligned myofibrils in wild-type myotubes differentiated with differentiation medium. In contrast, aberrant α-actinin-2 staining patterns were observed in heterozygous and homozygous myotubes. While heterozygous myotubes showed a less clearly defined cross-striation pattern due to an apparently distorted myofibrillar apparatus, homozygous myotubes displayed only an ill-defined myofibrillar α-actinin-2 staining (Fig. 3, left column). The latter defects could be fully restored by long-term electrical pulse stimulation, as indicated by a regular α-actinin-2 cross-striation staining pattern of highly ordered myofibrils as in the unstimulated wild-type (Fig. 3, right column). Long-term electrical pulse stimulation also appeared to affect the mitochondrial distribution, at least in homozygous myotubes, where the VDAC1 staining showed less coarse signals.

**Figure 1.**
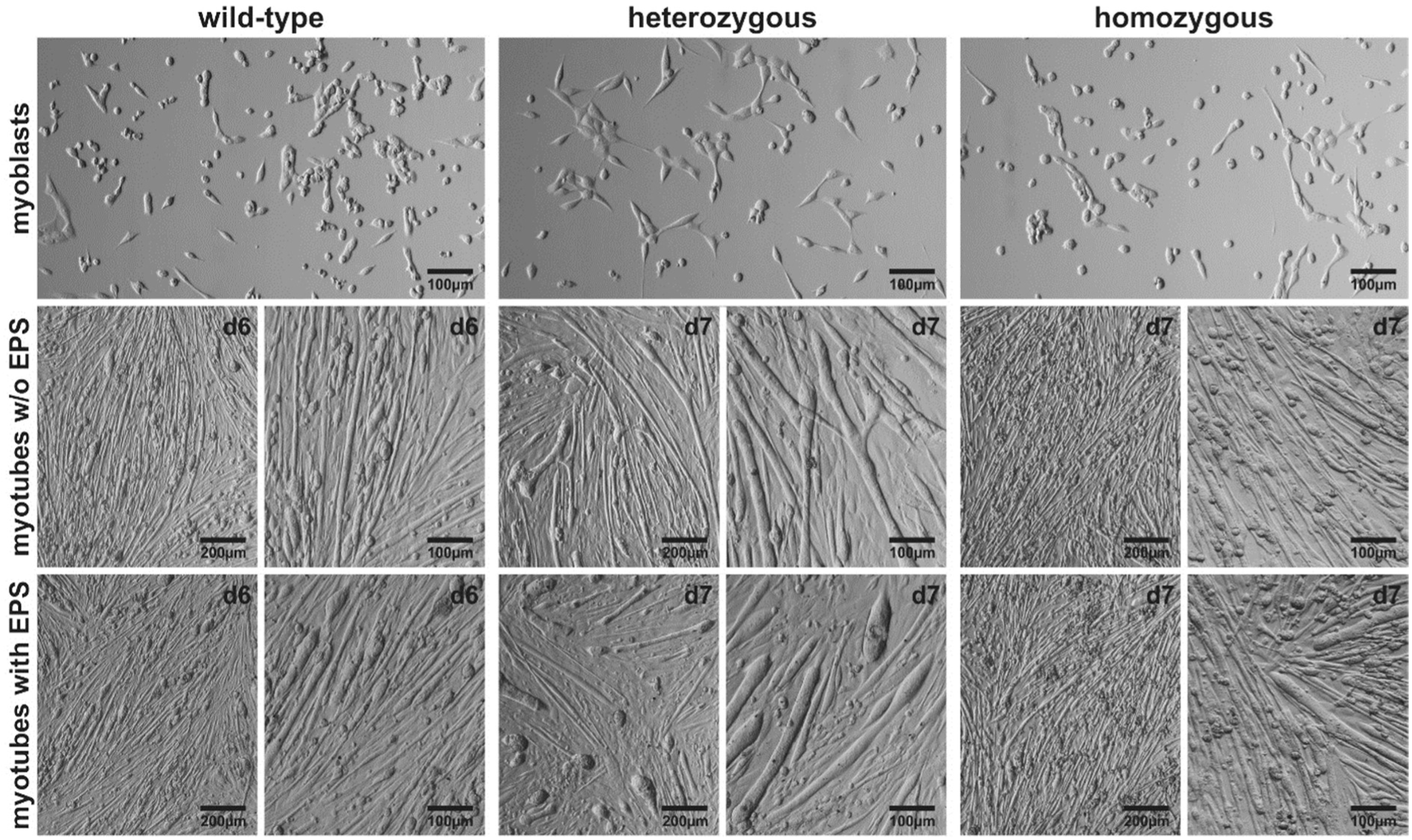
Formation of myotubes from immortalised murine myoblasts. Integrated modulation contrast (IMC) microscopy images of immortalised murine hetero-and homozygous R349P desminopathy and wild-type myoblasts and myotubes. Myotube formation in differentiation medium without or with electrical pulse stimulation (EPS) for six to seven days. Myoblast images, 100x magnification; myotube images, 50x and 100x magnifications. One representative polyclonal culture is shown for each genotype; wild-type (#224, gastrocnemius), heterozygous (#43, gastrocnemius), and homozygous (#34, soleus). On the light microscopy level, myotubes have formed in all myoblast cultures independent from EPS.

**Figure 2.**
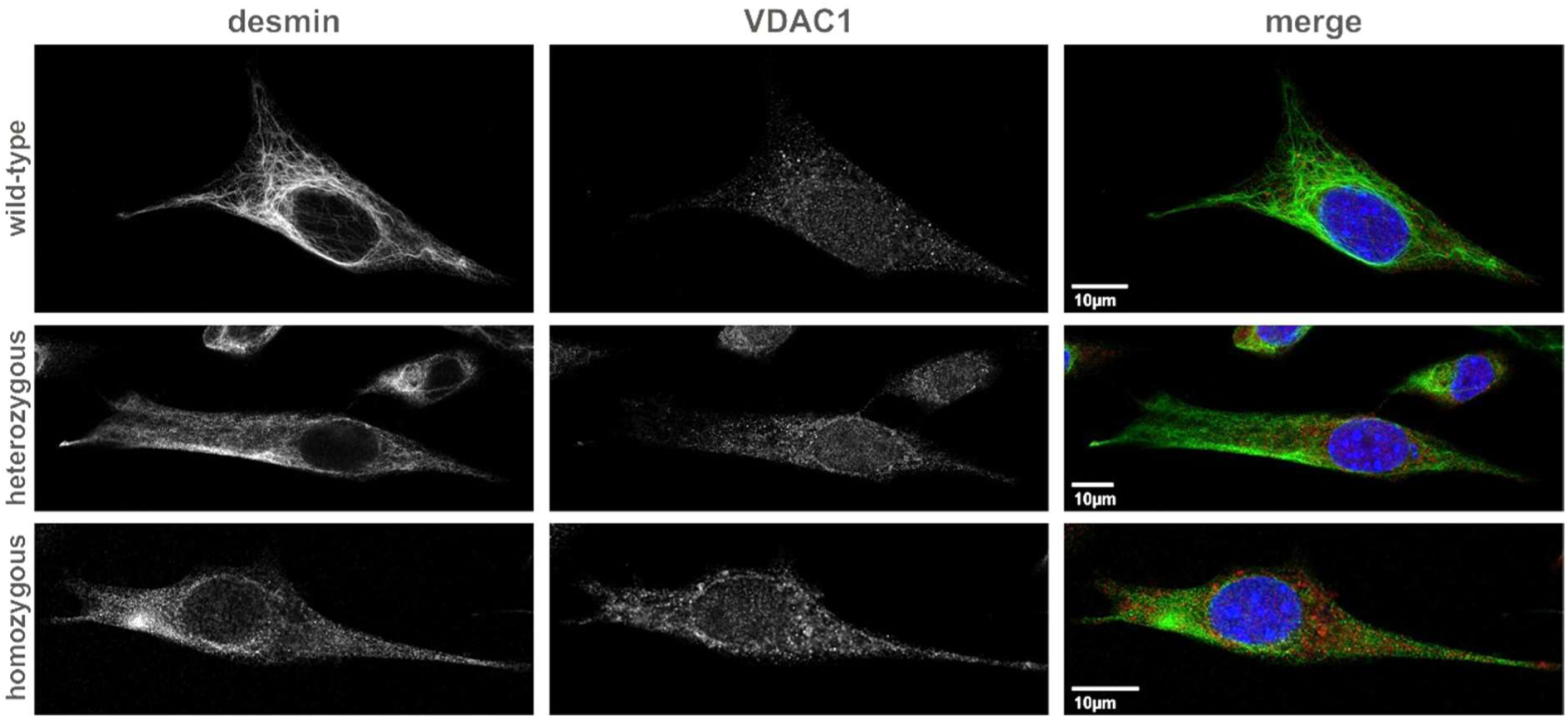
Impaired desmin intermediate filament network and mitochondria distribution in desminopathy myoblasts. Immortalised hetero-(#43, gastrocnemius) and homozygous (#34, soleus) R349P desminopathy as well as wild-type (#224, gastrocnemius) myoblasts were stained with antibodies directed against desmin (green), VDAC1 (red) and DAPI (blue). Confocal imaging showed an intact desmin intermediate filament network in wild-type myoblasts, and a collapsed network/desmin protein aggregation in homozygous cells. Heterozygous myoblasts displayed an intermediate phenotype with remnants of a filamentous network in conjunction with protein aggregates. The distribution of mitochondria in homozygous myoblasts had a coarser structure with an increased VDAC1 signal intensity. All images were deconvolved using Huygens Professional (Scientific Volume Imaging B.V., Hilversum, The Netherlands).

**Figure 3.**
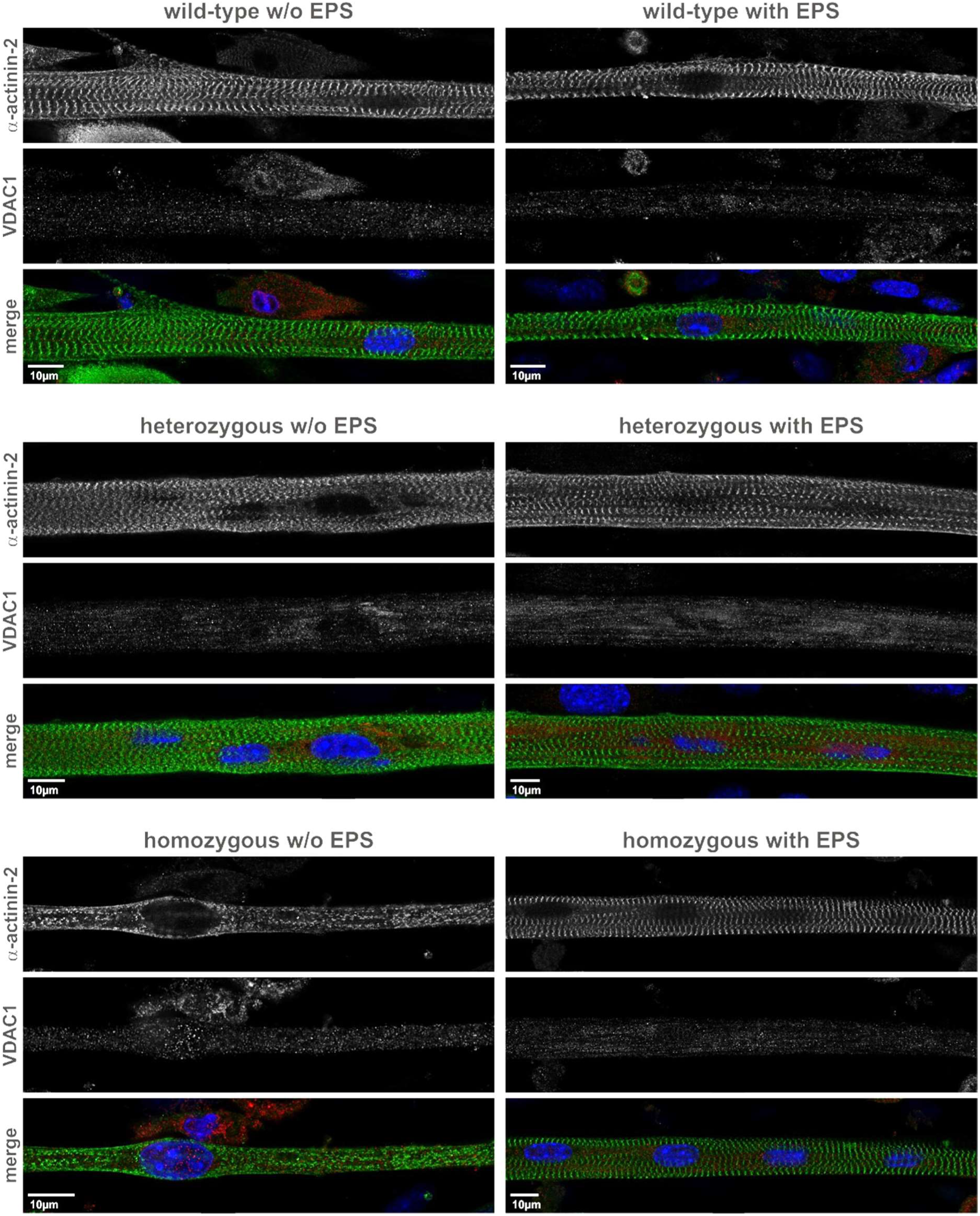
Electrical pulse stimulation strongly promotes the formation of myofibrils in desminopathy myotubes. Myotubes formed from immortalised hetero-(#43, gastrocnemius) and homozygous (#34, soleus) R349P desminopathy as well as wild-type (#224, gastrocnemius) myoblasts were stained with antibodies directed against α-actinin-2 (green), VDAC1 (red) and DAPI (blue). While confocal imaging showed myofibrils in wild-type myotubes, heterozygous and homozygous myotubes displayed a mostly unstructured distribution of α-actinin after six to seven days of differentiation without electrical pulse stimulation (EPS). The use of EPS during the differentiation period promoted the formation of myofibrils in the desminopathy myotubes to the level seen in wild-type cells and led to a less coarse distribution of mitochondria. All images were deconvolved using Huygens Professional (Scientific Volume Imaging B.V., Hilversum, The Netherlands).

### Significantly reduced proton leak in density gradient isolated mitochondria from desminopathy myotubes

We purified mitochondria from homozygous R349P desmin knock-in and wild-type myotubes using density gradients (Fig. 4A). Gradient fractions with a density of 1.14-1.16 g/mL containing mitochondria-compatible vesicular material and also both markers of mitochondrial matrix (citrate synthase) and membrane (VDAC1, TOMM20) were considered to contain structurally and functionally intact mitochondria (Fig. 4B,C). Aliquots of the freshly prepared mitochondrial suspensions were used for high-resolution respirometry focusing on the linear coupling controls (non-phosphorylating resting state, OXPHOS state, electron transfer capacity) with NADH-/complex I-linked (SUIT012, Fig. 4D) and succinate-/complex II-linked (SUIT020, Fig. 4E) substrates and the oligomycin-induced/complex V-inhibition non-phosphorylating resting state (SUIT020, Fig. 4E). The oxygen flux data obtained were normalised to the injected mitochondrial protein content. Repeated measurements of homozygous desminopathy and wild-type mitochondria showed no differences in respiration and, in particular, a normal function of complex I (Fig. 5A,B). Instead, the only statistically significant difference we detected was a reduced proton leak (L_Omy_, p=0.03) in the mitochondria of the R349P desmin knock-in myotubes (Fig. 5B).

**Figure 4.**
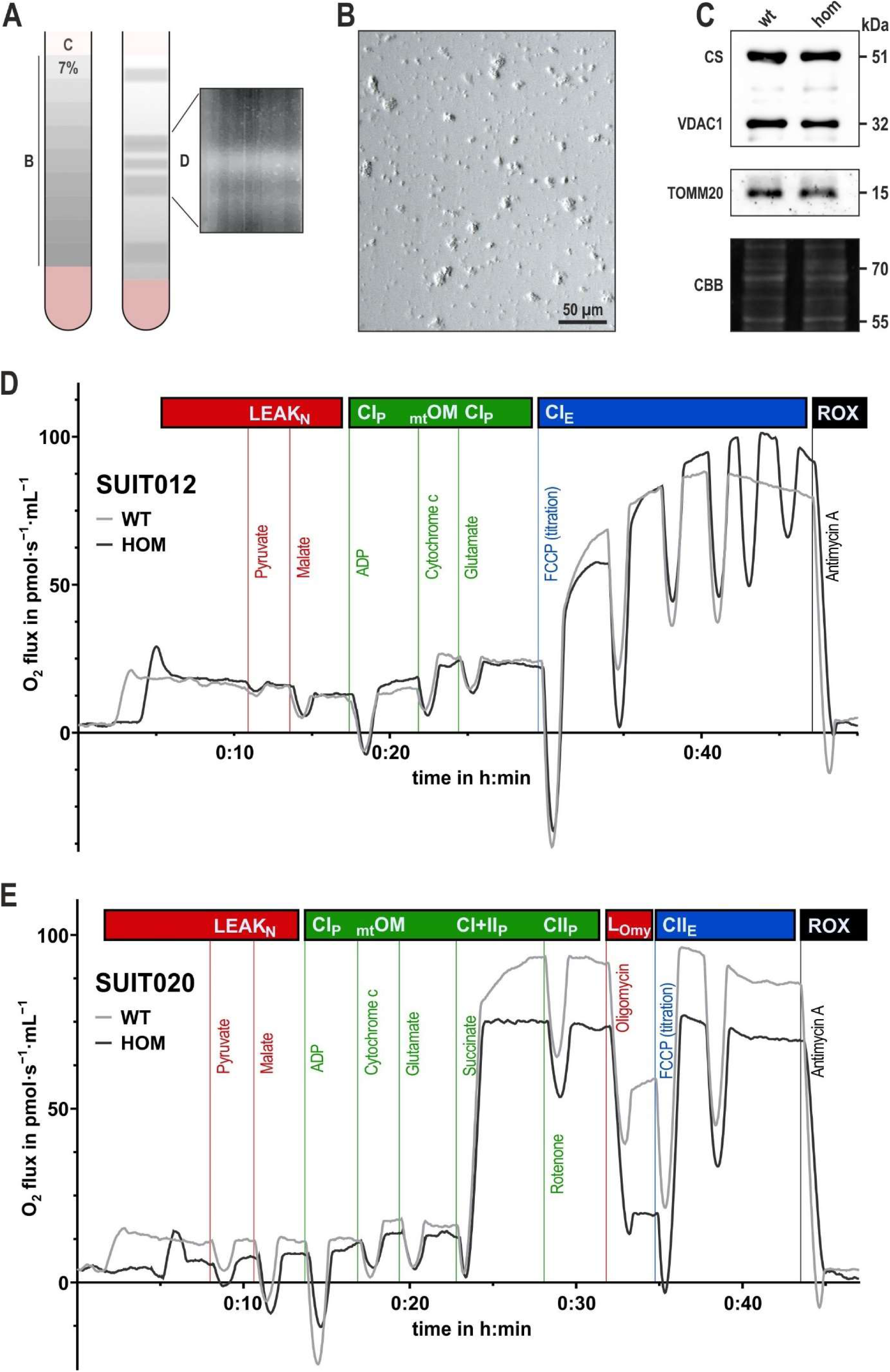
Isolation of structurally and functionally intact mitochondria from desminopathy and wild-type myotubes. (**A**) Myotubes formed from immortalised homozygous (#177, gastrocnemius) R349P desminopathy as well as wild-type (#224, gastrocnemius) myoblasts were harvested, lysed and used for density gradient isolation of mitochondria. A, crude mitochondrial preparation; B, gradient density steps; C, buffer overlay; D, after ultracentrifugation, fractions enriched in mitochondria were located in the center of the gradient. (**B**) Phase contrast microscopy image of a native smear of suspension from zone D of the gradient showing cellular material compatible in size and shape with mitochondria. (**C**) Immunoblotting using antibodies directed against citrate synthase (CS), voltage-dependent anion-selective channel 1 (VDAC1) and translocase of the outer mitochondrial membrane complex subunit 20 (TOMM20) of material derived from zone D of the gradient confirmed the presence of similar amounts of structurally intact wild-type and desminopathy mitochondria. CBB, fluorescence image of the Coomassie Brilliant Blue stained SDS-PA gel after blotting as a loading control. (**D**, **E**) Exemplary respirometry results showing the oxygen consumption over time. The curves shown have been normalised to the total protein content of the mitochondrial suspensions injected into the respirometer chambers. SUIT012, protocol assessing the linear coupling control (non-phosphorylating resting state, OXPHOS state, electron transfer capacity) with NADH-/complex I-linked substrates. SUIT020, protocol assessing i) the linear coupling control (non-phosphorylating resting state, OXPHOS state, electron transfer capacity) with focus on succinate-/complex II-linked substrates and ii) the oligomycin-induced/complex V-inhibition non-phosphorylating resting state. One out of a total of eight measurements per SUIT protocol is shown. LEAK_N_, non-phosphorylating resting state/respiration in the absence of adenylates at protocol start; CI_P_, NADH-linked substrate pathway in active OXPHOS state with saturating ADP; _mt_OM, cytochrome c control efficiency as a test for the integrity of the mtOM; CI+II_P_, NADH-and succinate-linked substrate pathways in active OXPHOS state with saturating ADP after further stimulation by addition of succinate; CII_P_, succinate-linked substrate pathway in active OXPHOS state with saturating ADP after inhibition of complex I with rotenone; L_Omy_, non-phosphorylating resting state/respiration after blocking complex V with oligomycin; CI_E_, uncoupler titration to obtain the electron transfer capacity for the NADH-linked substrate pathway; CII_E_, uncoupler titration to obtain the electron transfer capacity for the succinate-linked substrate pathway; ROX, baseline residual oxygen consumption due to oxidative side reactions after addition of complex III with antimycin A.

**Figure 5.**
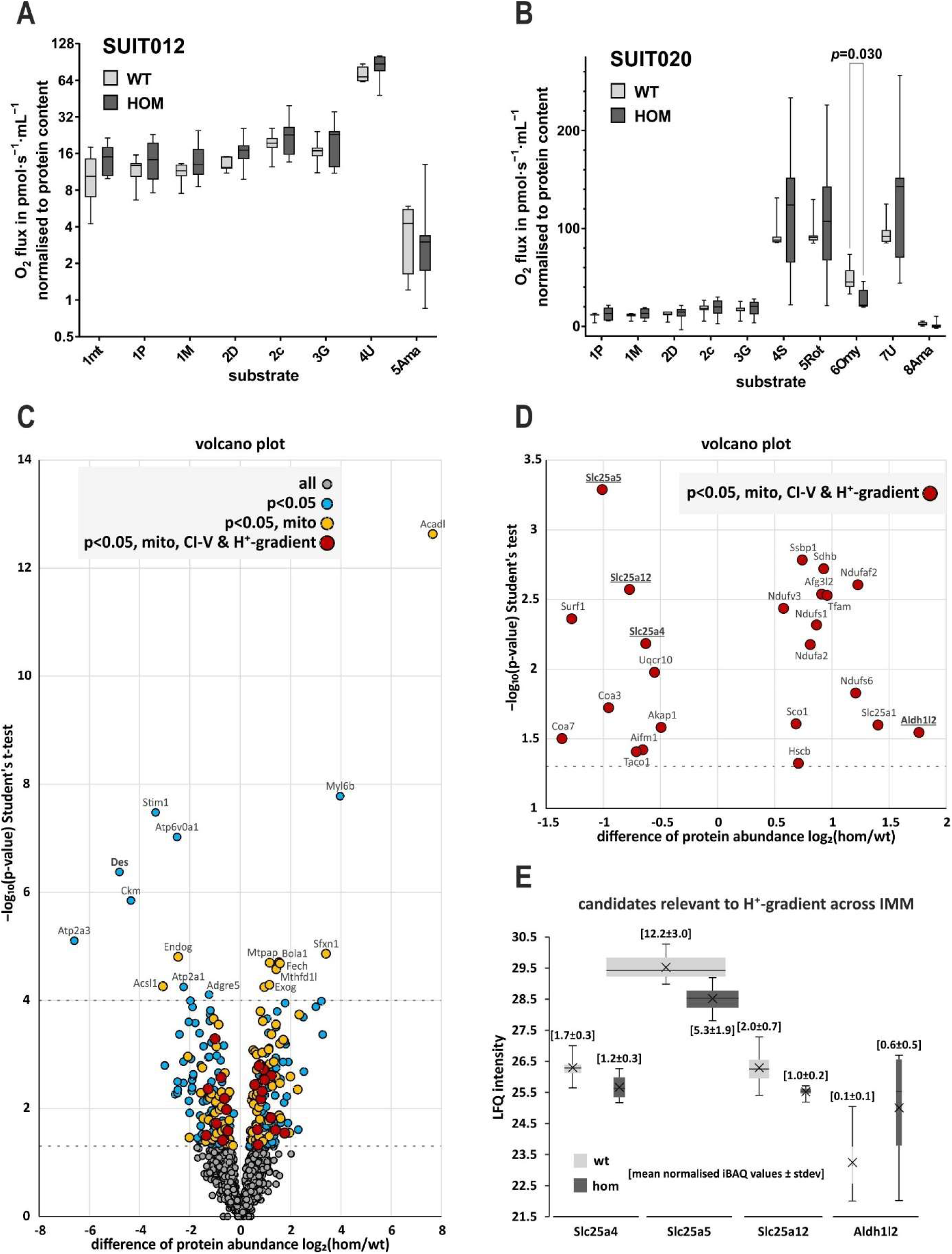
R349P desminopathy mitochondria display a significantly reduced proton leak. (**A**, **B**) Box plots of oxygen consumption normalised to the total protein amount, i.e., mitochondria amount measured for the different phases, divided into the different phases of substrate inductions and inhibitor applications during the SUIT012 and SUIT020 respirometry protocols. Boxes extent from the 25^th^ to 75^th^ percentiles, lines in the boxes indicate the medians and whiskers display min to max values from seven independent measurements per protocol; one replicate from each myoblast culture was identified as an outlier and excluded from further analysis. Statistical significance was calculated by the Mann-Whitney U test. 1mt, injected mitochondria; 1P, pyruvate addition; 1M, malate addition; 2D, ADP addition; 2c, cytochrome c addition; 3G, glutamate addition; 4U/7U, FCCP uncoupler addition; 4S, succinate addition; 5Ama/8Ama, antimycin addition; 5Rot, rotenone addition; 6Omy, oligomycin addition. (**C**, **D**) Volcano plots highlighting regulated proteins of homozygous versus wild-type mitochondria preparations, which were also used for injection into the respirometer. X-axis, log_2_- transformed mean fold change; y-axis, log_10_-transformed *p*-value. Grey circles, all proteins detected (1,270, out of which 505 (40%) were mitochondrial proteins); blue circles, subfraction of significantly regulated proteins with *p*≤0.05 (276), out of which 122 (44%) were mitochondrial proteins (orange circles); red circles, further subfraction of proteins related to mitochondrial complexes I to V and proton gradient across the inner mitochondrial membrane (enlarged view in (**D**)). Dotted lines indicate significance levels of 0.05 and 0.0001. (**E**) Box plots of the relative abundances across samples (log_2_- transformed LFQ intensities) of the four significantly regulated protein candidates involved in reducing the mitochondrial proton gradient across the inner mitochondrial membrane; additional crosses indicate mean values. Note that the horizontal box widths correspond to the mean normalised iBAQ values as a measure of the relative protein abundances within the same samples.

### The reduced proton flux across the inner mitochondrial membrane results from reduced levels of ADP/ATP translocases

The reduced proton leak (L_Omy_) in the presence of oligomycin-inhibited complex V in homozygous desminopathy mitochondria indicates either a decreased permeability of the inner mitochondrial membrane itself or a reduced, complex V-independent consumption of the mitochondrial proton gradient. To address the latter, we subjected aliquots of the same purified homozygous and wild-type mitochondrial fractions that were used for respirometry to label-free quantitative mass spectrometry. Analysis of the proteomic data revealed a total of 1,270 detected proteins, of which 505 were mitochondrial proteins (Fig. 5C, Tab. S1) based on an initial un-reviewed comparison with MitoCarta3.0 entries (https://www.broadinstitute.org/mitocarta/, (Rath et al., 2021)). Of the total of 1,270, 276 proteins were significantly (p≤0.05) regulated in the homozygous sample, of which a number of 122 proteins were mitochondrial proteins (Fig. 5C, Tab. S1) based on a manually verified comparison with MitoCarta3.0 and Uniprot database (https://www.uniprot.org) entries. A set of 52 mitochondrial proteins were downregulated in the homozygous mitochondria including two proteins related to complex I, one protein of complex III, and four assembly factors of complex IV, all seven of which are encoded by the nuclear DNA (Fig. 5D, Tab. S1). With respect to the detection of a reduced proton leak, three other proteins were identified whose carrier functions reduce the mitochondrial proton gradient by transporting protons into the mitochondrial matrix (Fig. 5D, Tab. S1>), namely Slc25a4 (ADP/ATP translocase 1, ratio hom/wt=0.65), Slc25a5 (ADP/ATP translocase 2, ratio hom/wt=0.50, with highest iBAQ value as indicated), and Slc25a12 (mitochondrial electrogenic aspartate/glutamate antiporter SLC25A12 or calcium-binding mitochondrial carrier protein Aralar1, ratio hom/wt=0.59). The set of proteins that were up-regulated in the homozygous mitochondria contained 70 entries. Among them, five proteins of and one protein related to complex I and one protein related to complex IV, all seven being encoded by nuclear DNA, and two proteins of complex II (Fig. 5D, Tab. S1). In addition, Aldh1l2 (mitochondrial 10-formyltetrahydrofolate dehydrogenase, ratio hom/wt=3.39, with lowest iBAQ value as indicated), an enzyme whose activity reduces the mitochondrial proton gradient by generating protons in the mitochondrial matrix, was upregulated (Fig. 5D, Tab. S1), as well as Slc25a1 (electroneutral mitochondrial tricarboxylate transport protein, ratio hom/wt=2.64). Furthermore, two proteins with roles in mtDNA maintenance, namely Ssbp1 (Single-stranded DNA-binding protein, ratio hom/wt=1.67) and Tfam (Transcription factor A, mitochondrial, ratio hom/wt=1.94) were upregulated (Tab. S1). A box plot illustrating the relative abundances across samples (log_2_-transformed LFQ intensities) in conjunction with the mean normalised iBAQ values as a measure of the relative protein abundances within the same samples (Fig. 5E) highlights ADP/ATP translocase 2 (Slc25a5), ADP/ATP translocase 1 (Slc25a4) and mitochondrial electrogenic aspartate/glutamate antiporter Slc25a12 as the most abundant and downregulated candidate proteins to account for the reduced proton leak. To corroborate the data at the tissue level, we re-evaluated proteomic data from the soleus muscle of heterozygous and homozygous R349P desmin knock-in mice and wild-type siblings generated for a previous publication (Winter et al., 2016). The protein levels of Slc25a4 were markedly reduced in heterozygous (ratio het/wt=0.47) and homozygous (ratio hom/wt=0.36) conditions as compared to the wild-type. Slc25a5 was also reduced in homozygous (ratio hom/wt=0.78) and almost absent in heterozygous (ratio het/wt=0.08) soleus muscle. Slc25a12 levels were reduced in both genotypes (ratios het/wt=0.81 and hom/wt=0.42), too.

## Discussion

In the present study, we newly generated immortalised myoblast cultures derived from skeletal muscle of heterozygous and homozygous R349P desminopathy mice and wild-type siblings, which could be well differentiated into spontaneously contacting myotubes. The myoblasts and myotubes reflect the previously described typical defects in the desmin intermediate filament network in muscle cells and tissues (Bär et al., 2005; Clemen et al., 2015). Our myoblast disease model was used to study the influence of electrical pulse stimulation-induced muscle cell contraction on the myofibrillar organisation and to further investigate the secondary mitochondrial damage in desminopathy.

### Electrical pulse stimulated contractions promote myofibril formation in desminopathy myotubes

A previous study focusing on the biomechanical properties of isolated skeletal muscle fibers from R349P desminopathy mice showed that mutant desmin disrupts the three-dimensional alignment and distorts the local angular axial orientation of myofibrils (Diermeier et al., 2017). These findings are further corroborated at the cellular level by the present work, as heterozygous myotubes showed impaired lateral alignment of myofibrils and homozygous myotubes showed only remnants of a myofibrillar cross-striation. Notably, long-term electrical pulse stimulation fully restored a normal myofibrillar pattern in both heterozygous and homozygous myotubes that was indistinguishable from the wild-type. The latter finding implies that long-term electrical stimulation of non-innervated myotubes over a period of one week is a new tool to improve the maturation of the cytoarchitecture of the myofibrillar apparatus *in vitro*. The molecular basis for this beneficial effect is currently unclear. The observation that patients deficient in desmin suffer from a myasthenic syndrome in addition to a primary myopathic disease (Durmus et al., 2016) highlights the negative impact of a defective extrasarcomeric cytoskeleton combined with impaired electrical stimulation of muscle cells due to a neuromuscular junction defect on the force-generating myofibrillar apparatus leading to early-onset generalised muscle weakness.

### Isolated desminopathy mitochondria show a reduced proton leak due to lower levels of ADP/ATP translocases

While a number of publications have shown the presence of mitochondrial defects in skeletal muscle from desminopathy patients and mouse models, only few have reported biochemical details. Analyses in muscle tissue homogenates derived from a heterozygous R350P desminopathy patient (Joshi et al., 2014) and heterozygous and homozygous R349P desminopathy mice (Winter et al., 2016) showed normal respiratory chain enzyme activities. Similarly, no change in mitochondrial respiration was detected in skeletal muscle homogenates derived from a heterozygous K240del desminopathy patient (Schröder et al., 2003, 2007). In contrast, the activities of complexes II and IV were decreased and the activity of complex I was lost in skeletal muscle homogenates from a heterozygous S13F desminopathy patient (McCormick et al., 2015). With regard to the K240del desminopathy, a more sophisticated analysis of single, saponin-permeabilized skeletal muscle fibers showed a reduced complex I activity (Schröder et al., 2003, 2007). Furthermore, analysis of the respiratory chain complexes in the soleus muscle of heterozygous and homozygous R349P desminopathy mice by native gel electrophoresis revealed reduced activities of complex I itself and as part of the supercomplex S_L_ (Winter et al., 2016). Thus, it appears that pathogenic desmin mutations may primarily affect the activity of mitochondrial respiratory complex I in skeletal muscle. However, the molecular pathophysiology including the initial steps leading to the observed defects is currently unclear. In this work, we used mitochondria purified from R349P desminopathy myotubes for high-resolution respirometry. Unexpectedly, we could not detect any significant changes in the activities of respiratory chain complexes I to IV. Instead, the only deviation we found was a significantly reduced proton leak in the R349P desminopathy mitochondria. Using quantitative proteomic analysis, we also found reduced levels of ADP/ATP translocases in the purified mitochondria, which could explain the observed reduced proton flux across the inner mitochondrial membrane. In addition, our re-evaluation of previously published proteomic data showed a reduction of ADP/ATP translocases in mature murine R349P desminopathy soleus muscle, along with the overall decrease in respiratory chain proteins (Winter et al., 2016).

Mitochondrial ADP/ATP translocases, also known as ADP/ATP carriers (AACs) or adenine nucleotide translocators (ANTs), which are members of the solute carrier family 25 of proteins (SLC25), are abundant proteins of the inner mitochondrial membrane that have two functions, ADP/ATP exchange and uncoupling by H^+^-influx. Apparently, the ADP/ATP exchange function competes to some extent with the H^+^-influx, but the ADP/ATP translocases also have a basal uncoupling proton flux (Bertholet et al., 2019). Notably, mitochondria isolated from mouse C2C12 myoblasts with a Slc25a4 (Aac1) and Slc25a5 (Aac2) double knock-out showed a strongly reduced basal respiration in the presence of oligomycin (Bertholet et al., 2019). This means that these mitochondria lacking two major ADP/ATP translocases also had a reduced proton leak, which fully supports our findings. The importance of ADP/ATP translocases is highlighted by the observation that mutations in *SLC25A4* (*ANT1*) cause myopathy and cardiomyopathy with multiple mtDNA deletions and increased mtDNA copy number in skeletal muscle (Echaniz-Laguna et al., 2012). Mitochondria with reduced levels of ADP/ATP translocases and consequent overload in the electron transport chain produce more reactive oxygen species (Wallace et al., 2010), leading to mtDNA damage and deletions (Beckman and Ames, 1998; Dos Santos et al., 2018; Wiesner et al., 2006). Large-scale mtDNA deletions have also been observed in the soleus muscle of homozygous R349P desminopathy mice (Winter et al., 2016). At the morphological level, these successive changes eventually result in enlarged mitochondria with widened cristae structures, as observed in skeletal muscle of R349P (Winter et al., 2016) and R405W (Batonnet-Pichon et al., manuscript in preparation) desminopathy mice. Thus, we consider our findings in purified mitochondria to be an early and important step in the pathogenesis of secondary mitochondriopathy in desminopathy.

## Declarations

## Authors’ contributions

C.S.C. conceived the study, designed experiments, and reviewed all data and statistical evaluations. C.B. and D.T. carried out experiments, developed protocols, analysed data, designed figures, and drafted the manuscript. D.P. provided respirometry measurement expertise and data analysis. D.T. performed statistical evaluations of respirometry values. B.E., K.M., and I.W. provided proteomic analyses. R.J.W. evaluated data and participated in writing the manuscript. C.B., R.S., and C.S.C. jointly finalised the manuscript text and figures. All authors have read and agreed to the final version of the manuscript.

## CRediT authorship contribution statement

**Carolin Berwanger**: Conceptualisation, methodology, investigation, validation, formal analysis, visualisation, writing - original draft, writing - review & editing. **Dominic Terres**: Conceptualisation, methodology, investigation, validation, formal analysis, visualisation, writing - original draft. **Dominik Pesta**: Methodology, resources, validation, formal analysis. **Britta Eggers**: Methodology, investigation, validation, formal analysis. **Katrin Marcus**: Conceptualisation, methodology, validation, formal analysis. **Ilka Wittig**: Methodology, investigation, validation, formal analysis. **Rudolf J. Wiesner**: Validation, resources, writing - review & editing. **Rolf Schröder**: Validation, resources, writing - review & editing. **Christoph S. Clemen**: Conceptualisation, methodology, resources, investigation, validation, formal analysis, visualisation, writing - original draft, writing - review & editing, supervision, project administration.

## Acknowledgements

We gratefully acknowledge the following support of this work, Peter F. M. van der Ven and Dieter O. Fürst, Institute for Cell Biology, University of Bonn, Bonn, Germany, for providing their electrical pulse stimulation expertise, Irmtrud Schrage-Knoll and Maria Bohmeier, Institute of Aerospace Medicine, German Aerospace Center, Cologne, Germany, for supporting the respiration measurements, and Patrick Lau, Institute of Aerospace Medicine, German Aerospace Center, Cologne, Germany, for general and organisational support during experimental work.

## Data availability

The data that support the findings of this study are available within the article and its supplementary materials.

## Supplementary materials

**Supplementary Table 1. Label-free quantitative mass spectrometry of total proteins from purified mitochondrial fractions from homozygous R349P desmin knock-in myotubes and wild-type cells.** Grey, all proteins detected; blue, significantly regulated proteins with p≤0.05; orange, significantly regulated mitochondrial proteins with p≤0.05; red, significantly regulated mitochondrial proteins with p≤0.05 related to mitochondrial complexes I to V and proton gradient across the inner mitochondrial membrane; bold and red, four significantly regulated protein candidates involved in reducing the mitochondrial proton gradient across the inner mitochondrial membrane. See also figure 5C-E. Raw data are available via ProteomeXchange with identifier PXD044624.

**Supplementary Table 2. Label-free quantitative mass spectrometry of total proteins from soleus muscle tissue of heterozygous and homozygous R349P desmin knock-in mice and wild-type siblings.** The full data set of the soleus muscle proteomic analysis from our previous study (Winter et al., 2016) has been re-assessed for the purposes of this work. Note that soleus muscles from five mice per genotype were pooled, and lysates prepared from these materials were subjected to label-free quantitative mass spectrometry analysis. Raw data are available via ProteomeXchange with identifier PXD046706.

